# Therapeutic doses of paracetamol with co-administration of cysteine and mannitol during early development result in long term behavioral changes in laboratory rats

**DOI:** 10.1101/2020.10.30.362137

**Authors:** Navneet Suda, Jasmine Cendejas Hernandez, John Poulton, John P Jones, Zacharoula Konsoula, Caroline Smith, William Parker

**Author notes:** Correspondence: William Parker, Ph.D., Department of Surgery, Duke University Medical Center, Box 2605.

## Abstract

Based on several lines of evidence, numerous investigators have suggested that paracetamol exposure during early development can induce neurological disorders. We had previously postulated that paracetamol exposure early in life, if combined with antioxidants that prevent accumulation of NAPQI, the toxic metabolite of paracetamol, might be innocuous. In this study, we administered paracetamol at or below the currently recommended therapeutic dose to male laboratory rat pups aged 4-10 days. The antioxidants cysteine and mannitol were included to prevent accumulation of NAPQI. In addition, animals were exposed to a cassette of common stress factors: an inflammatory diet, psychological stress, antibiotics, and mock infections using killed bacteria. At age 37-49 days, observation during introduction to a novel conspecific revealed increased rearing behavior, an asocial behavior, in animals treated with paracetamol plus antioxidants, regardless of their exposure to oxidative stress factors (2-way ANOVA; P < 0.0001). This observation would suggest that the initial hypothesis is incorrect, and that oxidative stress mediators do not entirely eliminate the effects of paracetamol on neurodevelopment. This study provides additional cause for caution when considering the use of paracetamol in the pediatric population, and provides evidence that the effects of paracetamol on neurodevelopment need to be considered both in the presence and in the absence of oxidative stress.

## Introduction

Several lines of evidence suggest that use of paracetamol in infants and young children may be detrimental to neurodevelopment [1]. To date, over a dozen studies, many well controlled for potentially confounding variables, have found links between neurodevelopmental disorders and paracetamol exposure early in life [2–15]. The initial indicator that a problem might exist was a 2008 survey which found that children who had a reaction to a vaccine and used paracetamol were more likely to have autism spectrum disorder (ASD) than children who had a reaction to a vaccine and did not use paracetamol [15].

Studies in laboratory mice and rats have demonstrated long lasting neurodevelopmental changes associated with exposure to one or two therapeutic doses of paracetamol during early development, when the brain is growing rapidly. Male animals are most susceptible to these changes, which occur far below the lethal dose of the drug, and include decreased speed of adaptation to a novel home environment in mice [16, 17], difficulties in learning in mice [16, 17], and biochemical changes in the brains of rats [18]. However, these studies generally use a slightly more than the maximum recommended paracetamol dosage in children of 14.7 mg/kg/dose. On the other hand, these studies involve fewer doses than the number received by human infants and children, who can be given up to 5 doses of paracetamol per day with a minimum waiting time between doses of 4 hours.

Of considerable concern is the interaction between oxidative stress and paracetamol. It is hypothesized that severe neurodevelopmental disorders can result from this combination [1]. Damage is thought to occur as a result of the production of the toxic metabolite of paracetamol, NAPQI, under conditions of oxidative stress. We have previously hypothesized that blockade of oxidative stress using glutathione precursors, known to be effective antidotes for paracetamol poisoning, might effectively eliminate the adverse effects of paracetamol on neurodevelopment [1].

In this study, we exposed laboratory rat pups to < 14.7 mg/kg/dose paracetamol formulated with the antioxidants cysteine, a glutathione precursor, and mannitol, a free radical scavenger. The ratio of paracetamol to cysteine and mannitol was maintained constant, using the ratio in FDA-approved intravenous paracetamol (OFIRMEV®). In addition, a cassette of factors that induce oxidative stress were utilized in the study. Factors known to induce oxidative stress were selected based on their common occurrence in the population, and included a Western diet [19], maternal stress [20, 21], infection [22], and antibiotics [23, 24]. Controls without paracetamol and without factors that induce oxidative stress were employed in a 2 x 2 study design. Adult male and female rats were preconditioned with a Western diet (high fat and high in processed sugar) prior to breeding, and maternal psychological stress was induced by bedding restriction and isolation during the first part of the third trimester of pregnancy. Exposure of male pups to a mock infection (killed lactobacillus), ampicillin, and paracetamol with antioxidants was carried out over a seven-day period using sub-cutaneous injections, from post-partum days 4 through 10 (P4 through P10). In this model, we hypothesized that neurodevelopment in animals without oxidative stress factors and with paracetamol plus antioxidants would be unaffected by paracetamol exposure. We also expected that neurodevelopment in animals with oxidative stress factors and with paracetamol plus antioxidants would be unaffected by the paracetamol exposure, assuming that the concentrations of antioxidants were sufficient. Although analysis of tissues could not be performed and behavioral tests were limited as a result of the impact of the emergence of SARS-CoV-2 on the study, sufficient data were obtained to undermine the hypothesis. Regardless of the presence of oxidative stressors, animals receiving paracetamol plus the antioxidants cysteine and mannitol on days P4 through P10 displayed substantial differences in their behavior on days P35 through P40 compared to controls.

## Methods

### Experimental design

All procedures were approved by the Duke Institutional Animal Care and Use Committee (Protocol A265-19-12). The design employed all oxidative stress inducing factors, both maternal and pediatric, as a single regimen. An outline of the experimental design and timeline devised with these considerations in mind is shown in **Figure 1**.

**Figure 1.**
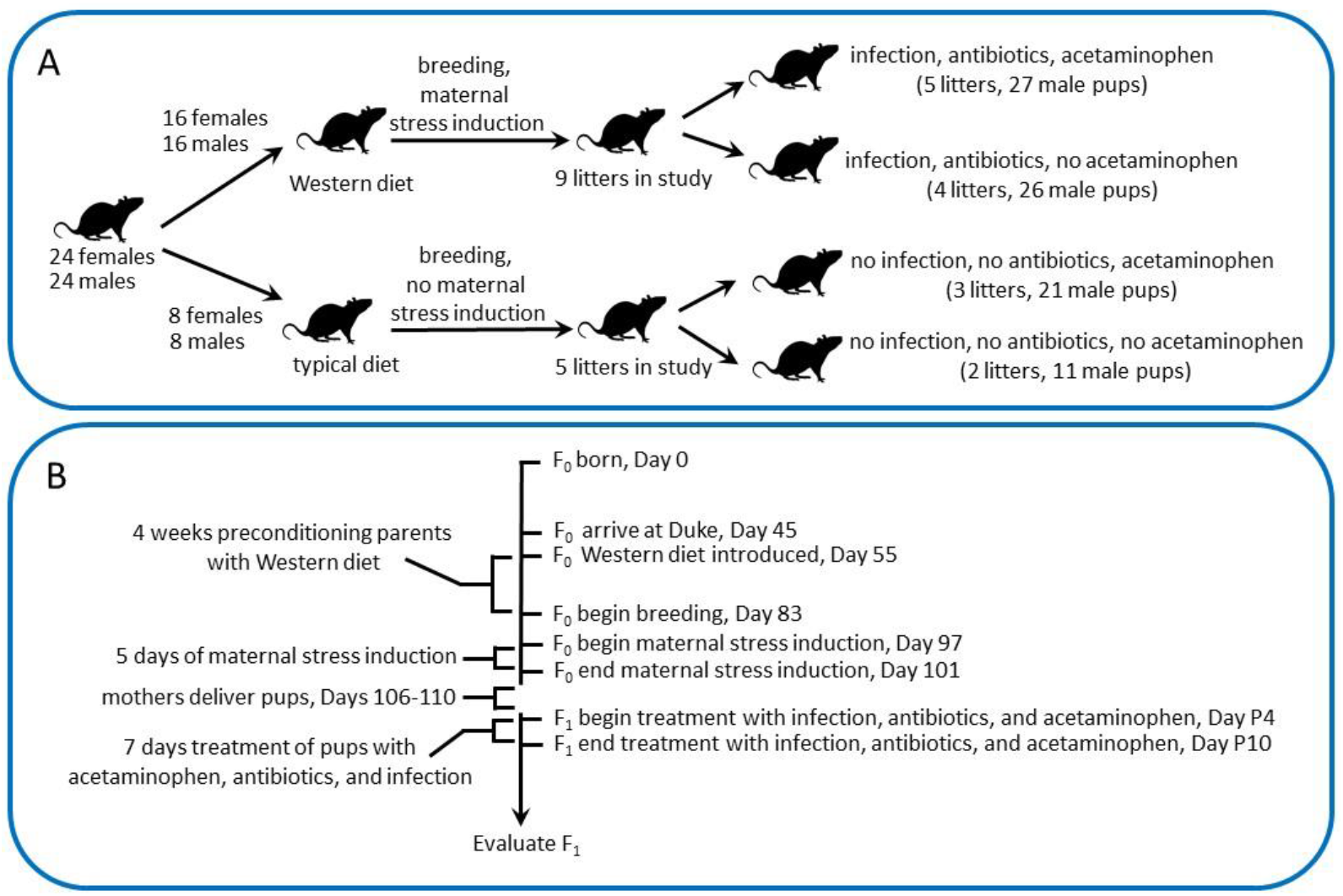
Experimental (A) design and (B) timeline. Rats were divided into four groups as described in the Methods. Two groups of animals received “oxidative stress induction” by preconditioning parents with a Western diet, maternal stress induction, and ampicillin treatment and mock infection with killed lactobacilli of male pups. Ampicillin and killed lactobacilli were administered together (mixed prior to injection) twice per day, 10 hours apart, for 7 days as described in the Methods. The other two groups received a normal laboratory rodent diet, no maternal stress induction, and injections with sterile saline as a control for injections with antibiotics and killed lactobacilli. Animals receiving oxidative stress induction were divided into two groups, with and without exposure to paracetamol. Similarly, animals receiving no oxidative stress induction were divided into two groups, with and with exposure to paracetamol. Paracetamol was administered every five hours for seven days as described in the Methods. All animals receiving no paracetamol received injections with sterile saline as a control.

Sprague Dawley rats (n = 24 male and n = 24 female) were purchased from Envigo. At the age of 55 days, 9 days after delivery to the animal care facility, rats were randomly assigned to an inflammatory “Western” diet (n = 16 male and n = 16 female), high in fat and processed sugar (Envigo rodent diet TD.88137) or retained (n = 8 male and n = 8 female) on a typical laboratory rodent diet (LabDiet® 5001 Rodent Diet). After 4 weeks of altered diet, rats were bred. After two weeks, the males were removed and female rats receiving a Western diet received induction of maternal stress by limiting bedding to approximately 2.1g of shredded paper (½ of a 2 cm thick by 8 cm in diameter disk of shredded paper; Bed-r’Nest®, The Andersons, Inc. Maumee, OH) from day E14 (approximately embryonic day 14) to the end of the day E18. Female rats receiving a typical diet, on the other hand, received extra enrichment and nesting material. Starting on the evening of day E19, animals were monitored every 5 hours around the clock for production of pups.

The pups of rats receiving maternal stress and a Western diet were randomly assigned to specific treatment arms. Pups were left with their natural birth dam rather than randomization for the following reasons: First, it was anticipated that some of the litter effects would take place in utero, and randomization at the time of birth could impede the possibility of their identification, providing potentially false positive results. Second, litters were born at different times, resulting in conspecifics that were not of comparable size. In addition, anticipated variation in pup weight imposed by psychological stress [25] was observed, and the difference in size of conspecifics was considered a potential risk factor for the smaller animals in the event of randomization. Finally, the additional stress from randomization of the pups was thought to be excessive in the face of the other stressors employed in this study. Given the design, particular attention was paid to potential influence of litter effects during evaluation of the data.

Starting on the morning of day P4 and ending at night on day P10, pups received subcutaneous injections of paracetamol, ampicillin, and mock infections with ampicillin-killed lactobacillus. Administration of paracetamol and ampicillin (described below) were adjusted for each pup based on the average weight of pups in that litter. Mock infections were not weight adjusted, and were held constant at a “volume equivalent” to 1 x 10^6^ bacteria per exposure (described below). Sterile saline was administered as a control (described below)

### Paracetamol

The maximum recommended exposure to paracetamol in infants is 14.7 mg/kg/dose with at least 4 hours between administrations and not more than 5 administrations in any given 24-hour period. Using these recommendations as limits for this study, pups were treated with paracetamol every 5 hours, starting in the morning of day P4 and ending at night on day P10. Paracetamol (OFIRMEV®, injection 1000mg/100ml) was appropriately diluted in sterile saline and injected sub-cutaneously in a total volume of 20 uL (for rat pups up to 13.5g), 30 uL (for rat pups up to 20g), 40 uL (for rat pups up to 27g), or 50 uL (for rat pups above 27g). The average weight of pups in each litter on days P3, P5, P7, and P9 was used to calculate the dilution necessary to achieve a 14.7 mg/kg/dose for each litter on the two subsequent days. For example, the average weight of each litter on day P3 was used to calculate dilutions for doses administered on days P4 and P5. Since the animals grew an average of 28% per day based on body weight, this approach insured that the dose administered, given errors in the amount injected (see above), did not exceed the recommended 14.7 mg/kg/dose. Further, as opposed to dose adjustment for each animal on a given day, this approach was technically feasible despite a protocol that required injection of more than 80 rat pups every five hours. Injection of sterile saline was used as a control.

The stock solution of paracetamol used (OFIRMEV®) contained 10 mg/ml paracetamol, 38.5 mg/ml mannitol, and 0.25 mg/ml cysteine hydrochloride, monohydrate. The stock solution was diluted in saline without altering the ratios of components.

Sub-cutaneous injections of 10 uL, 20 uL, 30 uL and 40 uL volumes were performed using a Vlow Medical (Netherlands) 3Dose™ Injector fitted with a 30 gauge by ½ inch hypodermic needle. Injectors were set at 10 uL injection units for all injections, and multiple units were dispensed depending on the volume needed. Injections using this approach had a standard deviation of less than 1 uL, or less than a 10% variance at 10 uL and less than a 5% variance at 20 uL. (Figure 2)

**Figure 2.**
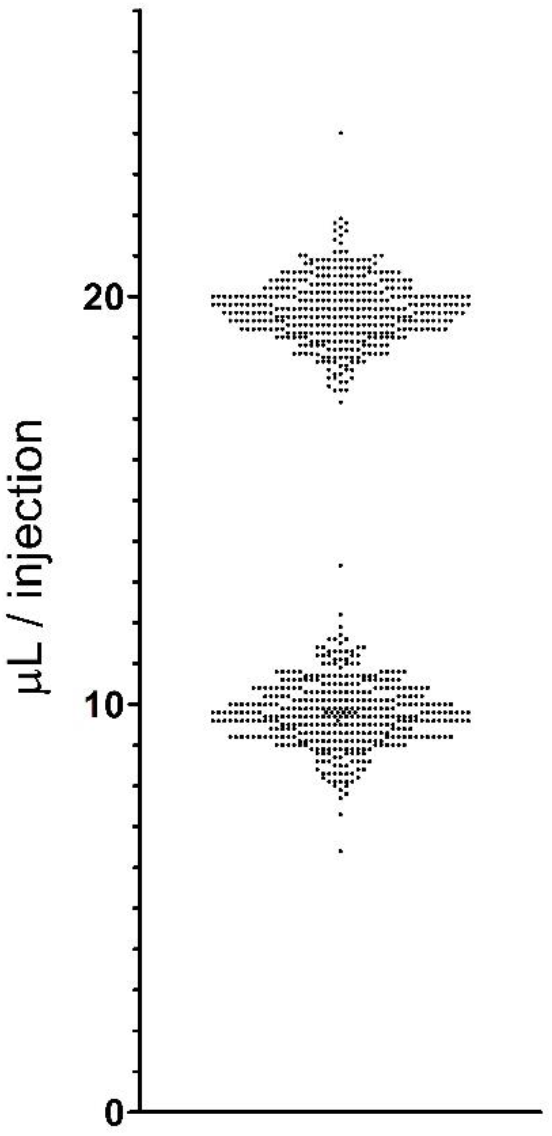
Precision of microinjections. The results of a total of 800 injections of sterile water with a Microdose ™ microinjector are shown. The injector was set at 10 uL for all injections, and volume was determined by weighing the solution using a Mettler balance and assuming a solution density of 1.00g/ml. One unit was dispensed for 400 injections, and two units (20 uL total) were dispensed for the other 400 injections. The single injections (expected volume of 10 uL) had a standard deviation of 0.83 uL with a coefficient of variation of 8.5%. The double injections (expected volume of 20 uL) had a standard deviation of 0.80 uL with a coefficient of variation of 4.1%.

### Ampicillin

Vials of ampicillin (Auromedics) were reconstituted with sterile water (≤ 1 EU/ml endotoxin; Sigma) according to the manufacturer’s instructions to obtain a buffered stock solution of 250mg/ml ampicillin. Stock solutions of ampicillin was flash frozen using liquid nitrogen and stored at −85 °C until the day of use. Rats were treated with appropriately diluted ampicillin twice per day, 10 hours between treatments, for 7 days (from P4 through P10). Ampicillin was administered sub-cutaneously in 10 uL of volume in a separate injection from paracetamol but at the same time as the paracetamol. Thus, all animals received four to five injections of paracetamol per day, with two of those injections coinciding with injections of 10 uL of ampicillin. Ampicillin was appropriately diluted to achieve a dose of 15 mg per kg of body weight per injection using average litter weights obtained one to two days before the injection, as described for paracetamol above. Ampicillin was combined with killed lactobacilli (see below) prior to injection. Injection of sterile saline was used as a control.

### Mock infections using ampicillin killed lactobacilli

Sub-cutaneious injection with a common, non-pathogenic bacterial strain killed by pre-treatment with ampicillin was utilized as a means of inducing a mock infection and subsequent inflammatory response with little to no danger of mortality in the study animals. For this purpose, cultures of *Lactobacillus reuteri* were be obtained from the ATCC (ATCC® 53608™, Lot 70021483) and grown in Criterion Lactobacilli MRS Broth (Hardy Diagnostics, Santa Maria, CA) at 37 °C under aerobic conditions with constant stirring for 48 hours. Bacteria were then incubated with 25 mg/L ampicillin for 1 hour at 37 °C with constant stirring. Next, ampicillin treated bacteria were harvested by centrifugation at 1536 x g for 10 minutes and washed 3 times with sterile saline. Finally, a 36.3% (volume/volume) slurry of ampicillin treated *Lactobacillus reuteri* in sterile saline was flash frozen and stored at −85°C until the day of use.

Macroscopic and microscopic examination of *Lactobacillus reuteri* revealed that the organisms were heavily aggregated, consistent with previous observations [26]. Because aggregation prevented accurate counts of individual bacterial cells, a volume of cell pellet equal to 1 x 10^6^ killed bacteria (with each bacteria estimated to have a volume of 0.85 um x 0.85 x 2.5 um = 1.8062 um^3^ [27]) was administered during each treatment. Killed bacteria were combined with ampicillin prior to administration and were delivered via subcutaneous injection twice per day with 10 hours between the two injections along with the ampicillin. Ampicillin and ampicillin killed lactobacilli were administered twice per day from P4 through P7, with 10 hours between each treatment. Sterile saline was administered as a control.

### Social Play Behavior

Social play behavior was assessed in rats between postnatal days 37 through 49 days in accordance with Veenema and colleagues [28]. Briefly, all subjects and stimulus animals will be handled for 5 consecutive days prior to the beginning of behavior testing. All experimental subject animals were isolated for 24 hours prior to behavioral testing to potentiate their motivation to engage in social interactions. All testing took place during the last two hours of the light phase. Animals were placed together with an unfamiliar animal belonging to the same experimental group. Clean cages with bedding were used, and activity was video recorded for 10 min. All behavioral videos were subsequently scored by an observer blind to the treatment condition of each animal using Solomon coder software (version beta 19.08.02; https://solomon.andraspeter.com/). Behavioral bins were as follows: exploration, play, investigation, chasing, rearing, auto grooming, and allogrooming. Play activity was further assigned to the following bins: nape attacks, boxing, pinning, and supine. Rearing was assessed for all animals (2 per cage), and all other behaviors were assessed for only one animal per cage. All videos had a total of 600.2 seconds of graded behavior.

### Statistics

Potential differences between experimental groups in weight and behavior were analyzed using unpaired t tests, 1-way ANOVA, or 2-way ANOVA, where appropriate. Comparisons between age and behavioral data were made using linear regression analyses. GraphPad Prism 8 software was used for all analyses.

## Results

### Effect of inflammatory factors on birth rate and pup survival

An overall summary of breeding results and assignment of litters to groups is shown in **Table 1**. Oxidative stress factors did not affect rates of pregnancy. However, 25% of the females with oxidative stress factors but not control females either abandoned their pups and/or demonstrated cannibalism.

**Table 1.** Animals and group assignments. ^a^One of the 6 pregnant animals without oxidative stress factors appeared pregnant initially but returned to a non-pregnant appearance without delivering any pups, presumably re-adsorbing her pups. ^b^Of the 12 pregnant females receiving oxidative stress factors, four of the females (25%) either abandoned their pups and/or demonstrated cannibalism. Of these, three were excluded from the study because they had no surviving male pups, and the fourth remained in the study with two surviving male pups. ^c^One male pup in the litter was sacrificed on Day P5 because it was in distress. ^d^One male pup in the litter was found dead on day P4 and another male pup in the litter was sacrificed on day P4 because it was in distress. All pups found dead or sacrificed due to distress were previously identified as runts within their litter.

| Oxidative stress factors | Number of animals | Number pregnant | Number male pups/litter completing study | Group assignment by litter |
| --- | --- | --- | --- | --- |
| No | 8 breeding pairs | 6<sup>a</sup> (25%) | 5,6,6,7,11 culled to 8 (32 total) | No APAP: 5, 6 (11 total) |
|  |  |  |  | APAP: 6, 7, 8 (21 total) |
| Yes | 16 breeding pairs | 12<sup>b</sup> (25%) | 2, 4, 5, 6, 6<sup>c</sup>, 6<sup>d</sup>, 8, 9 culled to 8, 10 culled to 8 (53 total) | No APAP: 6, 6, 6, 8 (26 total) |
|  |  |  |  | APAP: 2, 4, 5, 8, 8 (27 total) |

As shown in **Figure 3A**, the variation in pup weight/litter was substantially greater in litters born to mothers receiving a Western diet and psychological stress compared to mothers receiving normal diets and no stress. The range of pup weight/litter in animals receiving oxidative stress factors on day P3 was 4.56 g/pup/litter (n = 9 litters), about 14-fold greater than the range in litters without exposure to oxidative stress factors (range = 0.3211g/pup/litter, n = 5 litters).

**Figure 3.**
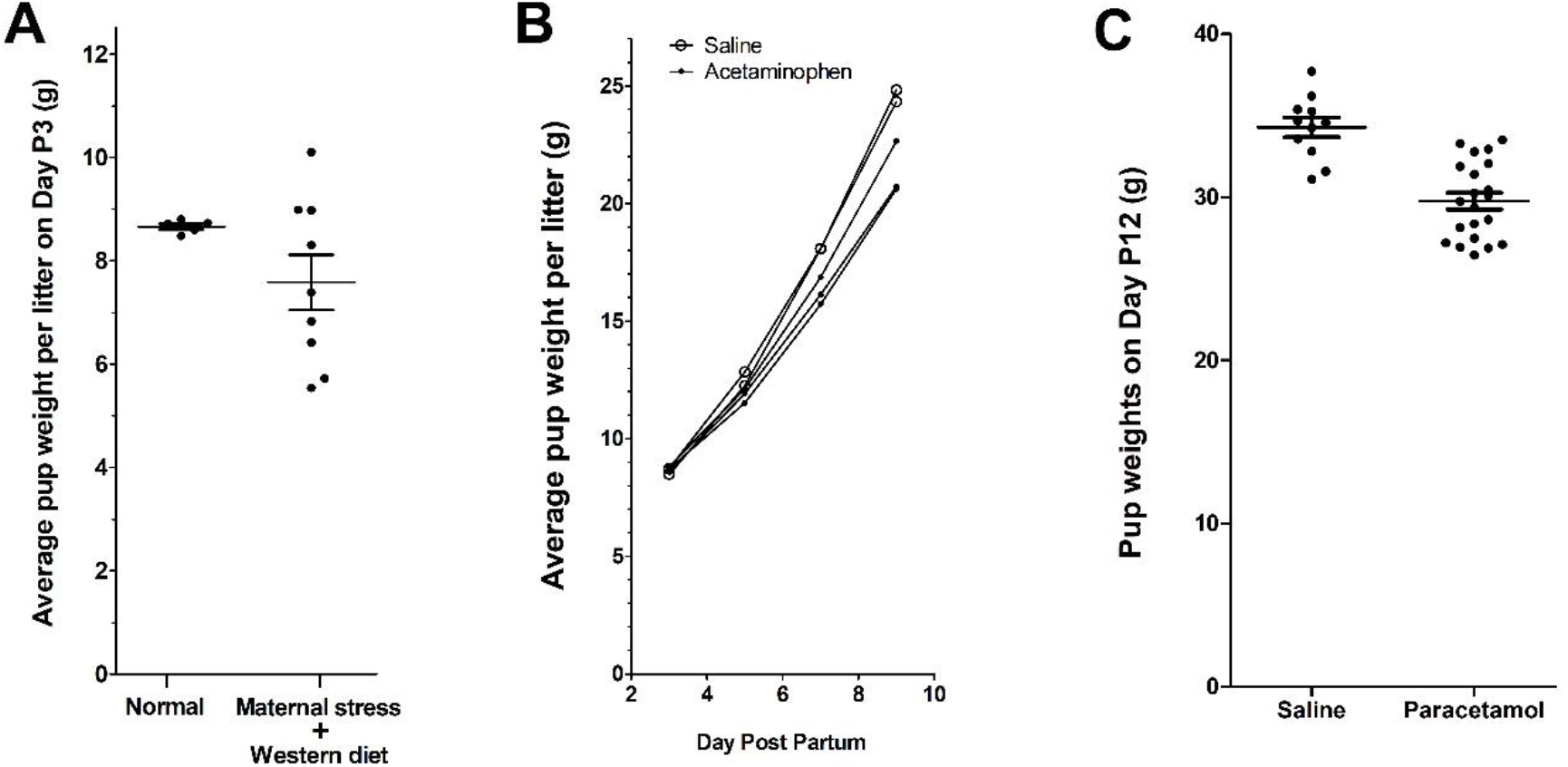
Weight of pups during the course of the experiment. In **Panel A**, the average weight of pups per litter on Day P3, prior to administration of paracetamol or saline, is shown. The average weights of pups in litters born to mothers with a Western diet and experimentally induced psychological stress were much more widely distributed (21.1% coefficient of variation) than were average weights of pups in litters born to animals without a Western diet and psychological stress (“Normal”; 1.5% coefficient of variation; F test: p = 0.0002). In **Panel B**, changes over time in the average pup weight per litter in animals without a Western diet and experimentally induced psychological stress are shown. **Panel C** shows the weights of individual pups born to animals without a Western diet or experimentally induced psychological stress as a function of paracetamol exposure versus saline as a control. Pups exposed to paracetamol weighed on average 13.2% less than pups receiving saline (t test: p < 0.0001), despite having essentially the same average weights on day 3 prior to initiation of drug exposure (Panel B).

Following the initiation of injections on Day P4, 32 out of 32 pups in the group receiving saline injections completed the study. Further, 56 out of 59 pups in the group receiving mock infections and antibiotics completed the study (Table 1).

### Effect of combined paracetamol plus antioxidants cysteine and mannitol

On day P3, prior to the initiation of injections, pups not receiving oxidative stress factors had very similar weights (Figure 3A). However, as shown in Figures 3B and 3C, weight gain was less in animals receiving paracetamol plus cysteine and mannitol than in controls receiving saline. This observation is consistent with the pharmaceutical labeling for injectable paracetamol (OFIRMEV®), which indicates that impaired growth as a result of drug administration in laboratory animals has been observed. This observation may be an effect of additives to the paracetamol preparation rather than to the paracetamol itself (see Discussion).

As shown in **Figure 4**, regardless of exposure to oxidative stress factors, treatment with paracetamol plus antioxidants (cysteine and mannitol) on days P4 through p10 was associated with a significant increase in an asocial behavior, rearing, while being exposed to a novel conspecific at age P37 through P49 (P < 0.0001). No significant association between rearing behavior and litter was observed in any experimental group (lowest p-value for any group = 0.56, average p value for all four groups = 0.73). Further, no significant effect of age on rearing behavior was observed in any group (lowest p-value for any group = 0.63, average p value for all four groups = 0.78), indicating that differences in behavior were due to the treatment rather than to litter effects or artifacts associated with the variable age of the animals. No statistically significant differences between any social behaviors (play or grooming) was observed associated with treatment with paracetamol or with exposure to oxidative stress factors.

**Figure 4.**
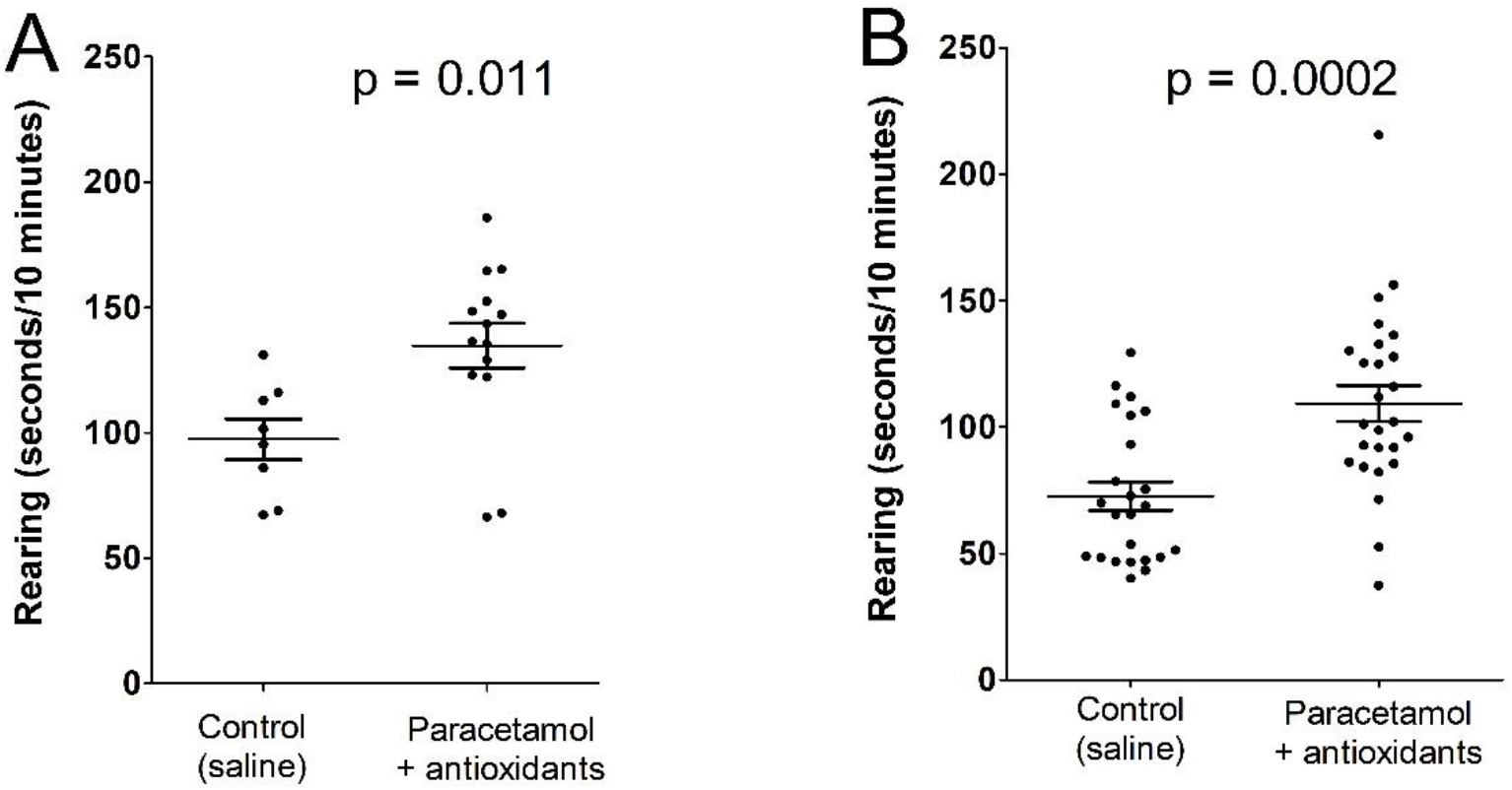
Effect of exposure to paracetamol plus antioxidants on days P4 through P10 on rearing behavior at age P37 through P49. Results are shown in **Panel A** for animals receiving no oxidative stress factors (see Methods), and in **Panel B** for animals receiving oxidative stress factors. Results of post-hoc t tests are shown. Analysis of the combined data by 2-way ANOVA showed significant effects of paracetamol + antioxidants (p< 0.0001) and of oxidative stress factors (p = 0.0036), but not for the interaction between the two (p = 0.97).

In animals not receiving oxidative stress factors, early life treatment with paracetamol plus cysteine was associated with 38.4% more rearing behavior at age P37 through P49 than observed in controls (p = 0.011). In animals receiving oxidative stress factors, treatment with paracetamol plus cysteine was associated with 50.6% more rearing behavior at age P37 through P49 than was observed in controls (p = 0.0002). However, assessment of rearing data from all four groups (with and without oxidative stress factors, with and without paracetamol plus cysteine exposure) using a 2-way ANOVA revealed no effect of interaction between exposure to oxidative stress factors and treatment with paracetamol (p = 0.97). This observation suggests that exposure to oxidative stress factors did not have a significant impact on the consequences of exposure to paracetamol plus cysteine in this model. The analysis by 2-way ANOVA confirmed a significant effect of treatment with paracetamol plus cysteine on rearing behavior (p < 0.0001). However, the analysis also revealed a significant effect of exposure to stress factors on rearing behavior under the conditions of this experiment (p = 0.0036). Interestingly, exposure to stress factors early in life was associated with *less* rearing behavior at age P37 through P49.

## Discussion

A wide range of observations from clinicians, immunologists, and epidemiologists suggest that exposure to paracetamol during early life, in combination with oxidative stress, can cause long-term neurological damage [1]. The mechanism by which this damage occurs apparently involves oxidative stress-induced accumulation of the toxic metabolite of paracetamol, NAPQI. Fortunately, a number of compounds, all metabolic precursors to the potent antioxidant glutathione, effectively aid in the reduction of NAPQI and reduce paracetamol toxicity [29]. Among these antidotes for paracetamol poisoning is cysteine, a component of the currently approved paracetamol formulated for IV use, OFIRMEV® (injection 1000mg/100ml).

We had previously postulated that inclusion of a glutathione precursor might eliminate the neurological damage imposed by early life exposure to paracetamol [1]. This study would suggest that this hypothesis is incorrect. One caveat to this conclusion might be that our experiments did not include sufficient precursor, in this case cysteine, and that additional precursor would have fully eliminated neurodevelopmental injury in this model. However, the inclusion of multiple oxidative stress inducers in the model did not significantly alter the outcome, suggesting that the neurodevelopmental effects we observed are, at least to some extent, independent of oxidative stress. Other caveats to this study need to be considered (see discussion below), but nevertheless it seems prudent at the present time to reject our hypothesis, establishing a new working model in which glutathione precursors, although effectively mitigating the gross effects of paracetamol overdose [29] or perhaps paracetamol exposure under conditions of oxidative stress, do not completely protect the developing brain from paracetamol-induced neurodevelopmental injury.

While the results presented herein might be considered alarming, this is not the first study demonstrating that exposure of laboratory animals to paracetamol during early development can cause long term neurological changes. For example, Viberg and colleagues at Uppsala University exposed mice aged P10 to two doses of paracetamol (30mg/kg/dose) four hours apart, and found an almost complete loss of their normal ability to learn how to navigate a radial arm maze when the animals were young adults, aged 2 months old [17]. Viberg also observed profound, long-term alterations in rearing behavior following the same paracetamol exposure [17]. The individual doses used by Viberg were more than twice as high as the doses used in children and in this study. However, current FDA guidelines indicate that drugs should be safe in laboratory animals at concentrations 6-fold higher than those used for testing in humans [30]. With this in mind, and given that Viberg only administered two doses, it is apparent that the currently approved use of paracetamol in infants and children would never have passed pre-clinical testing if the drug were evaluated based on current standards.

In this study, we did not test the effects of paracetamol under conditions in which oxidative stress was ensured. Although we utilized a novel “multi-dimensional” model to recapitulate the human experience that might lead to oxidative stress, we also included cysteine, a precursor for glutathione, along with all paracetamol treatments. In addition, the IV formulation we used contained copious amounts of mannitol, a free radical scavenger that probably has a positive effect on glutathione levels under conditions of stress [31]. Thus, this study does not address the hypothesis that severe adverse outcomes resulting from paracetamol-induced neurodevelopmental injury require oxidative stress, depletion of glutathione, and excess accumulation of NAPQI [1]. The need for testing of this hypothesis in a laboratory model is urgent. However, sufficient evidence from studies in humans already exists to warrant caution [1].

Limitations of this study include the fact that controls were not treated with mannitol and cysteine. Although both antioxidants are well tolerated, mannitol in particular might have potential effects on the biology of the system. For example, mannitol has been shown to lower hematocrit levels in laboratory rats [32]. Although speculative, mannitol may account for the decreased weight gain in the animals treated with paracetamol in this study. With these limitations in mind, additional studies are needed that include (a) controls with cysteine and mannitol, and (b) further behavioral testing of animals exposed the paracetamol with and without antioxidants.

## Acknowledgements

The authors wish to thank Uwe Scheuermann, Susan Poulton, Beth Weatherspoon, Susanne Meza-Keuthen, Randy Bollinger, Dawn E. Bowles, Zoie Holzknecht and Hazel Gelpi for their encouragement and support. The authors also thank April Kolstad, Eliott Mills, Jesse DeGraff, and Kaylee Lynn for their invaluable guidance and assistance with the animal studies. In addition, the authors thank Colin Duckett for facilitating completion of this project during the COVID-19 pandemic. Finally, we thank Staci Bilbo and Lauren Anderson for numerous helpful discussions and for their support.

## Notes

### Competing Interest Statement

The authors have declared no competing interest.

## References

1. Parker, W., et al., The role of oxidative stress, inflammation and paracetamol exposure from birth to early childhood in the induction of autism. Journal of International Medical Research, 2017. 45(2): p. 407–438.

2. Skovlund, E., et al., Language competence and communication skills in 3-year-old children after prenatal exposure to analgesic opioids. Pharmacoepidemiol Drug Saf, 2017. 26(6): p. 625–634.

3. Vlenterie, R., et al., Neurodevelopmental problems at 18 months among children exposed to paracetamol in utero: a propensity score matched cohort study. Int J Epidemiol, 2016. 45(6): p. 1998–2008.

4. Ystrom, E., et al., Prenatal Exposure to Paracetamol and Risk of ADHD. Pediatrics, 2017. 140(5).

5. Ji, Y., et al., Association of Cord Plasma Biomarkers of In Utero Paracetamol Exposure With Risk of Attention-Deficit/Hyperactivity Disorder and Autism Spectrum Disorder in Childhood. JAMA Psychiatry, 2019: p. 1–11.

6. Bittker, S.S. and K.R. Bell, Paracetamol, antibiotics, ear infection, breastfeeding, vitamin D drops, and autism: an epidemiological study. Neuropsychiatr Dis Treat, 2018. 14: p. 1399–1414.

7. Stergiakouli, E., A. Thapar, and G. Davey Smith, Association of Paracetamol Use During Pregnancy With Behavioral Problems in Childhood: Evidence Against Confounding. JAMA Pediatr, 2016. 170(10): p. 964–970.

8. Brandlistuen, R.E., et al., Prenatal paracetamol exposure and child neurodevelopment: a sibling-controlled cohort study. International Journal of Epidemiology, 2013. 42(6): p. 1702–1713.

9. Liew, Z., et al., Paracetamol use during pregnancy, behavioral problems, and hyperkinetic disorders. JAMA Pediatr, 2014. 168(4): p. 313–20.

10. Liew, Z., et al., Paracetamol use during pregnancy and attention and executive function in offspring at age 5 years. Int J Epidemiol, 2016. 45(6): p. 2009–2017.

11. Thompson, J.M., et al., Associations between paracetamol use during pregnancy and ADHD symptoms measured at ages 7 and 11 years. PLoS One, 2014. 9(9): p. e108210.

12. Liew, Z., et al., Prenatal Use of Paracetamol and Child IQ: A Danish Cohort Study. Epidemiology, 2016. 27(6): p. 912–8.

13. Liew, Z., et al., Maternal use of paracetamol during pregnancy and risk of autism spectrum disorders in childhood: A Danish national birth cohort study. Autism Res, 2016. 9(9): p. 951–8.

14. Avella-Garcia, C.B., et al., Paracetamol use in pregnancy and neurodevelopment: attention function and autism spectrum symptoms. Int J Epidemiol, 2016. 45(6): p. 1987–1996.

15. Schultz, S.T., et al., Paracetamol (paracetamol) use, measles-mumps-rubella vaccination, and autistic disorder. The results of a parent survey. Autism, 2008. 12(3): p. 293–307.

16. Philippot, G., et al., Adult neurobehavioral alterations in male and female mice following developmental exposure to paracetamol (paracetamol): characterization of a critical period. J Appl Toxicol, 2017. 37(10): p. 1174–1181.

17. Viberg, H., et al., Paracetamol (Paracetamol) Administration During Neonatal Brain Development Affects Cognitive Function and Alters Its Analgesic and Anxiolytic Response in Adult Male Mice. Toxicological Sciences, 2013. 138(1): p. 139–147.

18. Dean, S.L., et al., Prostaglandin E2 is an endogenous modulator of cerebellar development and complex behavior during a sensitive postnatal period. Eur J Neurosci, 2012. 35(8): p. 1218–29.

19. Auberval, N., et al., Metabolic and oxidative stress markers in Wistar rats after 2 months on a high-fat diet. Diabetology & metabolic syndrome, 2014. 6: p. 130–130.

20. Schiavone, S., et al., Severe life stress and oxidative stress in the brain: from animal models to human pathology. Antioxidants & redox signaling, 2013. 18(12): p. 1475–1490.

21. Eick, S.M., et al., Association between prenatal psychological stress and oxidative stress during pregnancy. Paediatric and perinatal epidemiology, 2018. 32(4): p. 318–326.

22. Novaes, R.D., A.L. Teixeira, and A.S. de Miranda, Oxidative Stress in Microbial Diseases: Pathogen, Host, and Therapeutics. Oxidative medicine and cellular longevity, 2019. 2019: p. 8159562–8159562.

23. El-Ansary, A., et al., The neurotoxic effects of ampicillin-associated gut bacterial imbalances compared to those of orally administered propionic acid in the etiology of persistent autistic features in rat pups: effects of various dietary regimens. Gut Pathog, 2015. 7: p. 7.

24. Kalghatgi, S., et al., Bactericidal antibiotics induce mitochondrial dysfunction and oxidative damage in Mammalian cells. Science translational medicine, 2013. 5(192): p. 192ra85–192ra85.

25. Salgado, S., M. Martínez, and C. Tarres, Body weight of litters of rats stressed during pregnancy. Medicina (B Aires), 1977. 37(1): p. 38–42.

26. MacKenzie, D.A., et al., Strain-specific diversity of mucus-binding proteins in the adhesion and aggregation properties of Lactobacillus reuteri. Microbiology, 2010. 156(11): p. 3368–3378.

27. Casas, I. and W. Dobrogosz, Validation of the Probiotic Concept: Lactobacillus reuteri Confers Broad-spectrum Protection against Disease in Humans and Animals. Microbial Ecology in Health & Disease, 2000. 12.

28. Veenema, A.H., R. Bredewold, and G.J. De Vries, Sex-specific modulation of juvenile social play by vasopressin. Psychoneuroendocrinology, 2013. 38(11): p. 2554–61.

29. Prescott, L., *Hepatotoxicity*, in *Paracetamol (Paracetamol) A Critical Bibliographic Review*. 1996, Taylor & Francis: Abingdon, UK. p. 319.

30. FDA, Guidance for Industry: Estimating the Maximum Safe Starting Dose in Initial Clinical Trials for Therapeutics in Adult Healthy Volunteers. 2005.

31. Shen, B., R.G. Jensen, and H.J. Bohnert, Mannitol Protects against Oxidation by Hydroxyl Radicals. Plant Physiol, 1997. 115(2): p. 527–532.

32. Kalemci, O., et al., Effects of Quercetin and Mannitol on Erythropoietin Levels in Rats Following Acute Severe Traumatic Brain Injury. Journal of Korean Neurosurgical Society, 2017. 60(3): p. 355–361.

